# An Automated and Robust Tool for Musculoskeletal and Finite Element Modeling of the Knee Joint

**DOI:** 10.1101/2023.10.14.562320

**Authors:** Amir Esrafilian, Shekhar S. Chandra, Anthony A. Gatti, Mikko Nissi, Anne-Mari Mustonen, Laura Säisänen, Jusa Reijonen, Petteri Nieminen, Petro Julkunen, Juha Töyräs, David J. Saxby, David G. Lloyd, Rami K. Korhonen

## Abstract

**Objective:** To develop and assess an automatic and robust knee musculoskeletal finite element (MSK-FE) modeling pipeline.

**Methods:** Magnetic resonance images (MRI) were used to train nnU-Net networks for auto-segmentation of knee bones (femur, tibia, patella, and fibula), cartilages (femur, tibia, and patella), menisci, and major knee ligaments. Two different MRI sequences were used to broaden applicability. Next, we created MSK-FE models of an unseen dataset using two MSK-FE modeling pipelines: template-based and auto-meshing. MSK models had personalized knee geometries with multi-degree-of-freedom elastic foundation contacts. FE models used fibril-reinforced poroviscoelastic swelling material models for cartilages and menisci.

**Results:** Volumes of knee bones, cartilages, and menisci did not significantly differ (*p*>0.05) across MRI sequences. MSK models estimated secondary knee kinematics during passive knee flexion tests consistent with *in vivo* and simulation-based values from the literature. Between the template-based and auto-meshing FE models, estimated cartilage mechanics often differed significantly (*p*<0.05), though differences were <15% (considering peaks during walking), i.e., <1.5 MPa for maximum principal stress, <1 percentage point for collagen fibril strain, and <3 percentage points for maximum shear strain.

**Conclusion:** The template-based modeling provided a more rapid and robust tool than the auto-meshing approach, while the estimated knee biomechanics were comparable. Nonetheless, the auto-meshing approach might provide more accurate estimates in subjects with distinct knee irregularities, e.g., cartilage lesions.

**Significance:** The MSK-FE modeling tool provides a rapid, easy-to-use, and robust approach to investigating task- and person-specific mechanical responses of the knee cartilage and menisci, holding significant promise, e.g., in personalized rehabilitation planning.

## I. Introduction

MUSCULOSKELETAL (MSK) disorders are major contributors to global disability. Knee osteoarthritis (KOA) is one of the most prevalent MSK disorders, affecting approximately 23% of individuals aged over 40 [1], with different phenotypes, such as mechanical overload and inflammation [2], [3]. Therefore, to ameliorate KOA, investigating pathophysiological mechanisms underlying its onset and progression requires a comprehensive and personalized understanding of the mechanobiological environment of the knee. However, acquiring *in vivo* knee mechanics involves highly invasive procedures [4], [5], which are infeasible in most cases. Moreover, such *in vivo* data is restricted to joint-level quantities, whereas tissue-level biomechanics (e.g., stress and strain) have been shown to govern cartilage degradation [6]–[8]. Therefore, developing non-invasive methods to obtain tissue-level knee biomechanics is crucial for advancing our understanding and improved KOA management.

Recent research has demonstrated the potential of computational modeling, namely musculoskeletal (MSK) and finite element (FE) models, to estimate personalized knee joint mechanobiological environment and degradation response [6], [9]–[12]. Indeed, FE models have been widely used to estimate the knee’s within tissue biomechanics (e.g., cartilage stress and strain) governing tissue degradation response [6]–[8], [12], [13]. Furthermore, MSK models have traditionally been used to estimate knee joint-level biomechanics to provide subject- and task-specific loading and kinematics boundary conditions for FE models to estimate within tissue biomechanics [14]. However, creating personalized MSK and FE models is time-consuming and labor-intensive, requiring several weeks of high-level expertise [15]. The process involves image segmentation, processing of the segmented geometries, meshing, model assembly, material model incorporation, estimation and incorporation of loading and boundary conditions, and achieving a converged solution [16], [17]. This need for specialized knowledge presents a barrier to broader use and limits reproducibility [18] of MSK and FE models in research and clinical settings where personalized models could have a significant impact on health outcomes [9], [11], [19], [20].

Several studies have developed rapid and (semi-)automatic methods to create knee joint models. Template-based methods have enabled rapid generation of knee MSK-FE models [17], [21] with the potential to estimate KOA progression [21] or plan personalized rehabilitation [9]. However, these methods are not fully automated, e.g., they require manual measurement of knee anatomical dimensions from medical images. Fully automated knee FE modeling algorithms have been developed based on auto-segmentation of magnetic resonance imaging (MRI) of subjects’ knees [22], [23]. Nonetheless, these offer no potential to produce the subject’s MSK model [22], [23], are limited to a particular MRI sequence [22], [23], exclude some knee tissues such as menisci [22], can take hours to create the subject’s FE model [23], and more importantly, often require manual preprocessing of the medical images [23]. Alterations in inputs to the FE model (e.g., obtained from different MSK modeling frameworks) [14] and excluding tissues such as the menisci [24] can substantially alter the FE model’s estimated knee biomechanics. To the authors’ knowledge, the existing automatic knee modeling approaches use rigid or linear elastic material models for knee cartilage and menisci [22], [23], which can largely affect the estimated knee biomechanics [25]. Notably, a nonlinear composite material model is essential to simultaneously assess mechanical responses of both fibrillar (collagen) and nonfibrillar (proteoglycans) matrices within the cartilage and menisci [6], [25], [26]. Also, it is necessary to include poro(visco)elastic material model to replicate fluid-flow-related mechanisms of porous tissues, as up to 85% of a dynamic load is carried by within-tissue fluid pressurization [27], [28].

In this study, we developed and assessed a robust and fully automated knee joint segmentation and MSK and FE modeling tool. The tool performs preprocessing of medical images and auto-segmentation of knee bones, cartilages, menisci, and major knee ligaments, and creates and assembles MSK and FE models of the subject’s knee. To improve the robustness of the tool, we developed two modeling pipelines, consisting of 1) template-based and 2) auto-meshing. To this end, we leveraged previously developed and validated template-based [17], [21] as well as hexahedral meshing [22], [29] algorithms. We compared knee biomechanics estimated by the two MSK and FE modeling pipelines and two different MRI sequences, as well as against a manually created FE model.

## II. Method

The modeling tool developed in this study (Fig. 1 and supplementary materials, Fig. S1) creates personalized MSK and FE models based on auto-segmentation of the subject’s MRI. Following the auto-segmentation, the tool generates two distinct sets of MSK and FE models, i.e., template-based and auto-meshing models. While the personalized MSK and FE models could be run independently, they were generated based on the same geometries and knee coordinate systems, enabling integrated sequentially-linked MSK-FE modeling [14]. We implemented the tool in Python and partly in MATLAB (using MATLAB engine for Python), with a graphical user interface (GUI, Fig. S1).

**Fig. 1.**
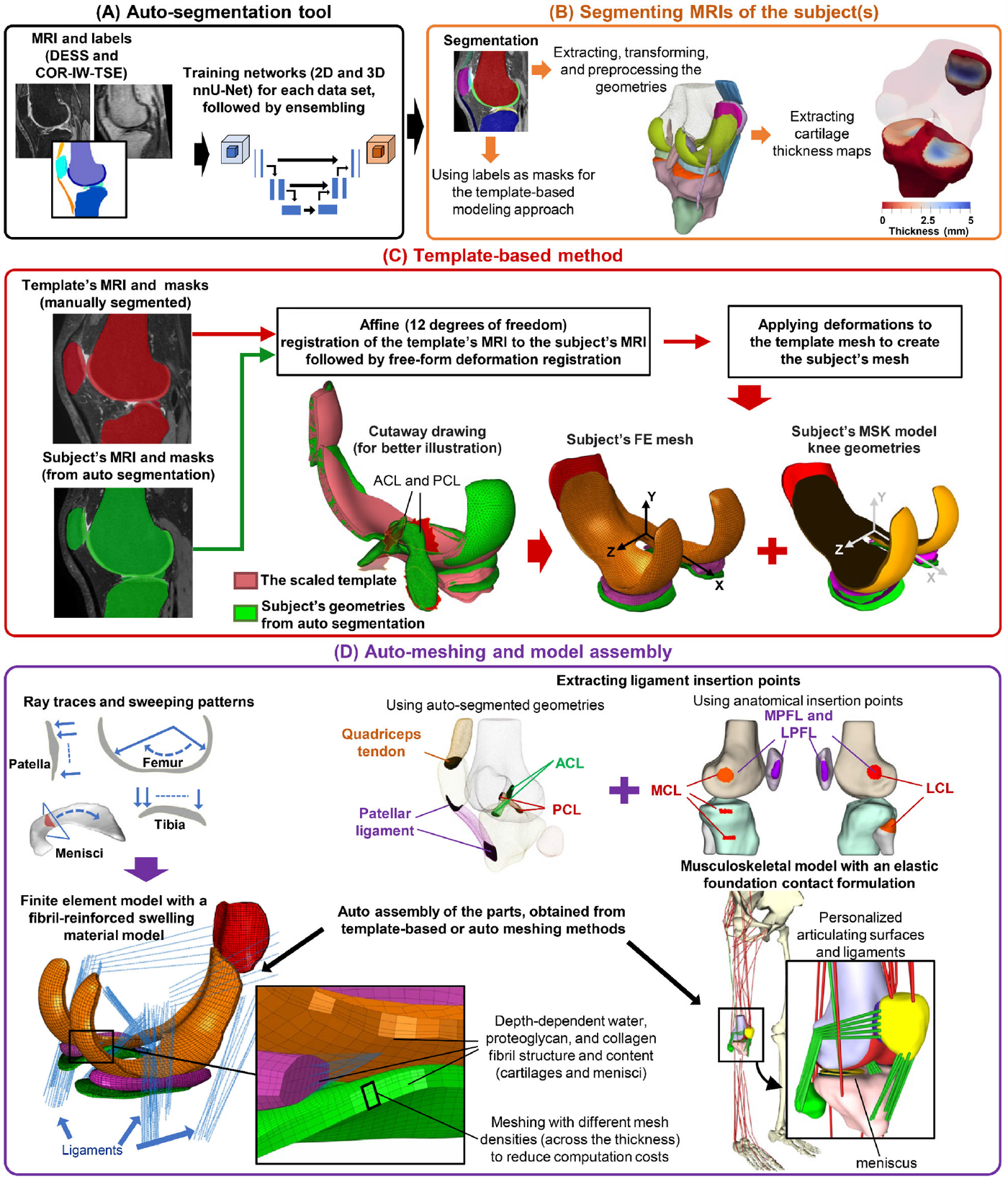
The segmentation and modeling pipeline developed in the study.

The MSK modeling exploited a previously developed and validated MSK model of the lower extremity [30]–[32], implemented in OpenSim v4.4 [33], [34]. The knee in the MSK model had 24 degrees of freedom (DoF), including the tibiofemoral joint, patellofemoral joint, medial meniscus, and lateral meniscus, each possessing 6 DoF [31]. The knee was constrained by twelve bundles of nonlinear (quadratic) elastic ligaments and capsular structures, as well as menisci horn attachments (Fig. 1) [35], [35], [36]. Knee contacts were a nonlinear elastic foundation model [37], [38].

The FE models included femoral, tibial, and patellar cartilages, menisci, and major knee ligaments. The knee ligaments were represented as bundles of nonlinear elastic springs, similar to the MSK model. The FE model’s ligament bundles consisted of anterior and posterior cruciate ligaments (ACL and PCL), medial and lateral collateral ligaments (MCL and LCL), medial and lateral patellofemoral ligaments (LPFL and MPFL), and patellar ligament. Within the FE model, cartilages and menisci were modeled using a fibril-reinforced poroviscoelastic swelling (FRPVES) material model [39].

We conducted comparisons of knee biomechanics among the MSK and FE models generated through the template-based and auto-meshing methods, as well as against a manually created FE model. More details are presented in the following subsections and supplementary materials.

### A. Subjects’ MRI data sources

MRIs from four datasets were used to train and assess the tool (Fig. S2). Three publicly available datasets, totaling 669 knee joints, were used to train the auto-segmentation tool. The first dataset was a subset of the Osteoarthritis Initiative (OAI) dataset and had knee MRIs from 507 knee joints. Segmentations for distal femoral and proximal tibial bones and femoral and tibial cartilages were provided by the Zuse Institute Berlin (ZIB) [40]. We used dual-echo steady-state (DESS) images (in-plane resolution of 0.3646 mm, slice thickness of 0.7 mm) and coronal intermediate-weighted turbo spin-echo (COR-IW-TSE) images (in-plane resolution of 0.3646 mm, slice thickness of 3 mm) [41]. The second dataset was the Stanford Knee MRI with Multi-Task Evaluation (SKM-TEA) [42] with 155 knee MRIs. DESS images were provided (in-plane resolution of 0.3125 mm, slice thickness of 0.8 mm) along with manual segmentations for femoral, tibial, and patellar cartilages and menisci. The third dataset was from the Open Knee project subjects (OKS) with seven knee MRIs [43]. The 3D fat-suppressed T_1_-weighted (FS-T_1_, DESS approximate) images were provided (in-plane resolution of 0.3516 mm, slice thickness of 0.7 mm) along with manual segmentations for bones (distal femur, proximal tibia and fibula, and patella), cartilages (femur, tibia, and patella), menisci, quadriceps tendon, patellar ligament, ACL, PCL, MCL, and LCL.

The fourth dataset, collected at the Kuopio University Hospital (Philips Achieva 3.0T scanner, Philips, Eindhoven, The Netherlands), was used to assess the MSK and FE modeling tool. The data collection was approved by the ethical committee of the Kuopio University Hospital (#140/2017) in accordance with the Helsinki Declaration. All participants provided written consent. The dataset consisted of MRIs from 52 knee joints (both knees of 26 subjects, ∼60% female), of which 19 subjects were healthy, and 7 subjects had medial tibiofemoral KOA (Kellgren-Lawrence [44] grade >1, with evident thinning of medial tibiofemoral joint space). We used proton density-weighted volume isotropic turbo spin echo acquisition with spectral attenuated inversion recovery (PDW-VISTA-SPAIR, DESS approximate) images with an in-plane resolution of 0.3571 mm and a slice thickness of 0.5 mm, and Proton Density (COR-IW-TSE approximate) images with an plane resolution of 0.303 mm and a slice thickness of 0.52 mm (Fig. S2). We reported the modeling results from PDW-VISTA-SPAIR images in the manuscript, while the results from Proton Density images are in the supplementary material.

### B. Auto-segmentation

We adopted 2D and 3D nnU-Net [45], a previously developed algorithm, to perform auto-segmentation (Fig.1-A and B). Before training, ITK-SNAP [46] and NiftyReg [47] libraries were used to reorient all the MRIs to LPI orientation (i.e., left-to-right, posterior-to-anterior, and inferior-to-superior), followed by resampling to 160×384×384 slices (as in OAI dataset). Next, we trained three different sets of networks, as follows in sections II.B.1 and II.B.2. We trained both 2D and 3D nnU-Net with five-fold cross-validation [45]. The default setting provided by the nnU-Net tool was used, except for the batch and patch sizes, which were increased to use the total capacity of an NVIDIA RTX A6000 for training. The 2D network had a batch size of 80 and a patch size of 384×384, while the 3D network had a batch size of 2 and a patch size of 96×256×256. Input images to the networks were augmented, consisting of mirroring, rotation (30°), isotropic scaling (50%), randomized Gaussian noise, Gaussian blur, brightness, contrast, gamma correction, and simulation of low resolution [45].

#### 1) DESS segmentation

Validation studies have shown DESS sequence provides accurate information on cartilage morphology (e.g., volume and thickness) [48], [49]. Thus, we used DESS as reference sequence in this study. Specifically for the OKS dataset, we used a more aggressive data augmentation consisting of non-rigid deformation, in which we registered the OKS dataset to the whole ZIB dataset using 12 DoF affine transformations (NiftyReg tool [47]). After the training, the best configuration was obtained using an ensemble algorithm [45].

#### 2) COR-IW-TSE segmentation

A considerable number of clinical datasets include MRI sequences other than DESS. To increase the applicability of the modeling tool developed in this study, we trained networks using COR-IW-TSE sequence, as well. In this, we first segmented bones, ligaments, and menisci for the ZIB dataset using the network trained in section II.B.1. Then, we checked all the labels, with more focus on cartilages, menisci, and bones, to ensure the accuracy of the segmentation. Lastly, COR-IW-TSE images of the ZIB dataset (507 knees) were used to train 2D and 3D nnU-Net networks and obtain the best ensemble [45], as in section II.B.1.

### C. Reconstruction of tissue geometry and preprocessing

The trained networks (section II.B) were used to auto-segment the fourth dataset of the study, consisting of 52 knee joints and two MRI sequences. Femoral and tibial bones and cartilages were used from the ZIB network, menisci and patellar cartilage from the SKM-TEA network, and fibula and patella bones, ligaments, and quadriceps tendon from the OKS network.

Following auto-segmentation, the marching cubes algorithm [50] via Visualization Toolkit Python library (VTK v9.2) [51] was used to extract the triangular surface mesh of each tissue. The extracted geometries were re-meshed with 1 mm edge size for bones and cartilages and 0.5 mm edge size for ligaments using the MeshLab Python library [52], [53]. Next, geometries underwent smoothing by a discrete Gaussian filter (variance = 0.15, ITK Python library) [54], Taubin filter (MeshLab Python library) [52], [55], and three iterations of Laplacian smoothing (MeshLab Python library) [52], [56]. We assumed each bone consisted of a singular connected volume and, thus, all enclosed volumes, except the largest one, were regarded as segmentation artifacts and subsequently excluded. Last, the knee coordinate system was determined, and all the surface meshes were saved in stereolithography (STL) file format (Fig. 1-B). Details on determining the knee coordinate system are provided in the supplementary material (section S1.1).

We also extracted femoral, tibial, and patellar cartilage thickness maps (Fig. 1-B), using VTK Python libraries [51], [57] by casting rays from bone vertices in their normal directions. If the ray had two intersections with the corresponding cartilage geometry (within an arbitrarily-chosen range of ±1 cm), the distance between the two intersections was considered cartilage thickness, and its magnitude was assigned to that vertex on the bone. Consequently, the cartilage thickness map had a resolution equal to the number of bone vertices (Fig. 1-B). We then calculated the average cartilage thickness for the load-bearing regions of medial and lateral femoral, tibial, and patellar cartilages (Figs. 2 and S3), with details provided in supplementary materials (section S1.2). Thickness maps were extracted prior to applying the smoothing filters.

**Fig. 2.**
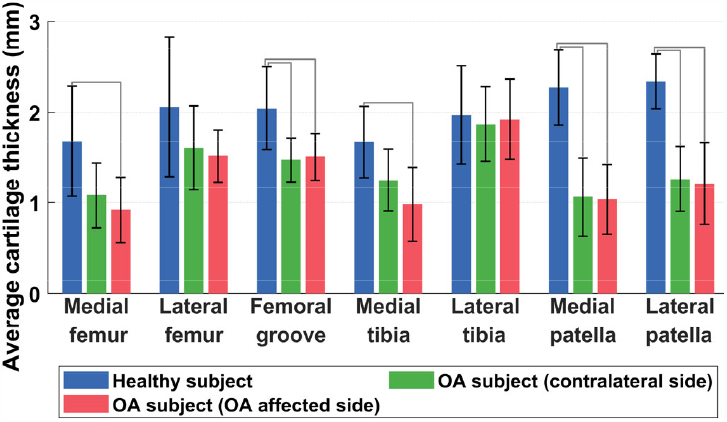
Knee cartilage thickness obtained from auto-segmentation of PDW-VISTA-SPAIR (DESS approximate) MRI of the fourth dataset of the study. Gray lines indicate statistically different results (*p*<0.05).

### D. Template-based MSK and FE modeling

We developed the template-based MSK and FE modeling pipeline to ensure the tool consistently produced a personalized model without failure. We adopted our previously developed and validated template-based modeling [9], [17], [21], in which a template model was scaled according to the anatomical dimensions of the subject’s knee, such as mediolateral and anteroposterior knee dimensions. In this current study, we used the same template MRI and FE model (with the FRPVES material model incorporated) but modified the scaling pipeline [23] to automatically transform the template MRI to the subject’s MRI. The template was from a healthy 33-year-old male and consisted of femoral, tibial, and patellar cartilages, menisci, ACL, PCL, LCL, MCL, LPFL, MPFL, patellar ligament, and quadriceps tendon [17]. The use of one template (rather than choosing between several templates [23]) has shown more promising results in predicting KOA progression and classifying subjects into correct KOA grade groups [21].

To provide a fully automated template-based modeling [23], we used a non-rigid Free-Form Deformation (FFD) algorithm [5] to obtain a displacement field that transforms the template MRI to the subject’s MRI. In this, tissue labels from auto-segmentation (section II.B) were used as masks (i.e., regions of interest). To enhance the FFD algorithm, the template MRI was first transformed into the subject’s MRI using a 12 DoF non-rigid affine registration [23], [47]. NiftyReg was used to conduct FFD and affine registrations [47], [58]. Using NiBabel in Python [59], the affine transformation matrix and FFD displacement field were then used to morph the template FE mesh (i.e., nodal coordinates) and create personalized FE geometries.

Ultimately, the final personalized FE meshes were converted to surface meshes, i.e., in STL format, to create the template-based MSK model. The contact algorithm within the MSK model [38] necessitated shell-type contact surfaces rather than 3D water-tight volumes. Thus, the cartilage-subchondral surface of the morphed template was considered the bone surface, while the opposing surface was considered the cartilage contact surface within the knee joint of the template-based MSK model (Fig. 1-C). The top and bottom surfaces of the menisci obtained from the morphed FE template were used as menisci geometries within the MSK model (Fig. 1-C).

### E. Auto-meshing MSK and FE modeling

#### 1) The hexahedral meshing algorithm and FE model

We adopted a previously developed hexahedral meshing algorithm [22], [29] to create a structured mesh for the femoral, tibial, and patellar cartilages and menisci from the auto-segmented geometries (Fig. 1-D). The algorithm used a swept set of point origins and raytracing techniques to construct rectangular grids [22], [29]. However, we slightly modified the sweeping functions (without altering the total mesh volume) to mesh the cartilages and menisci according to the tissue structure and composition while considering the FE modeling bottlenecks and computation costs. Mesh density (i.e., element size) was set according to mesh sensitivity analyses from our previous studies [6], [17], [25].

For the femoral cartilage, a mesh with an element surface area of approximately 1 mm^2^ and one element through the thickness was first created, as developed by Rodriguez-Vila et al. [29]. The mesh was then split into four layers across the thickness, representing Benninghoff arcades of collagen fiber orientation (Fig. 1-D) [60]–[62]. For the tibial and patellar cartilages, we updated the sweeping algorithm developed by Rodriguez-Vila et al. [29], which used radial sweeping to produce a mesh that is fine at the center of the cartilage but progressively coarser towards the edges (Fig. S4-A, also Fig. 4 in [29] and Fig. 6 in [22]). This can potentially result in contact instability and an increased number of elements for a suitably fine mesh, leading to substantial computational costs. Here, we used sweeping in Cartesian coordinates rather than in radial direction to create a computationally efficient mesh (Figs. 1-D and S4-B) [63]. In this, the algorithm first finds the mean (equivalent to vector sum) of all the surface normals and then uses this vector to create a grid (1×1 mm) and sweep the tibia and patellar cartilages (Fig. 1-D). The points from sweeping were then used to create the hexahedral mesh, with one element through the depth. Next, the mesh was split into three layers across the thickness. Last, each element on the contact surface was split into four elements [63] (Figs. 1-D and S4-B).

A circumferential sweeping algorithm for the menisci was used as developed by Rodriguez-Vila et al. (Figs. 1-D and S4-C) [29], from which we extracted the menisci element nodes. However, exploiting our previous knee FE modeling studies [6], [17], [25], we rearranged the nodes to create a more structured mesh (i.e., aspect ratio ∼1 and corner angle ∼90°) with four layers in the distal-proximal direction (Fig. S4-D). Following the meshing of cartilages and menisci, node and element sets, and contact surfaces required for assigning the material model and implementing loading and boundary conditions were generated automatically.

#### 2) The auto-meshing MSK model

In the MSK model, bones (i.e., distal femur, proximal tibia, and patella) and cartilages (i.e., femur, tibia, and patella) were used from each subject’s auto-segmented geometries (section II.C). To ensure compatibility with the required contact algorithm [38], which necessitated shell-type contact surfaces, we used a process of casting rays from each triangle of the bone surface toward its associated cartilage tissue. By identifying the point of farthest intersection between the ray and the cartilage, we obtained the shell-type cartilage contact surfaces (Fig. 1-D). Meniscal surfaces were obtained by converting the auto-generated hexa-hedral mesh (section II.E.1) to STL format and selecting the top and bottom surfaces, similar to the template-based MSK model.

### F. Ligaments and quadriceps tendon

All the MSK and FE models, either template-based or auto-meshing approach, had identical ligament insertion points, stiffness, and slack length. The insertion points of the ACL and PCL on the femoral and tibial bones, as well as the insertion points of the patellar ligament and quadriceps tendon on the tibial and patellar bones, were determined using auto-segmented geometries and the nearest-neighbor concept. In this, the points on the femur, tibia, and patellar bones with less than 2 mm distance from each end of the aforementioned ligaments (from auto-segmentation) were considered insertion points, corresponding to distal and proximal ends. The auto-segmentation of LCL and MCL was relatively poor (Table 1); thus, the insertion points of LCL, MCL, LPFL, and MPFL were obtained from the segmented bones, according to the literature (Fig. 1-D) [32], [64]–[67]. If the algorithm failed to detect any ligament insertion ends (including ACL and PCL), the corresponding insertion points were used from the morphed template model. We used this approach to enhance modeling robustness, particularly in cases of suboptimal ligament segmentation. The insertion points of the remaining ligaments, e.g., popliteofibular ligament and posteriomedial capsule within the MSK model, were untouched. The slack length and stiffness of all the ligaments were set according to Lenhart et al. [30], and menisci horn attachments according to Villegas et al. [68]. More details on the ligaments’ insertion point detection, material model, and parameters are provided in the supplementary material (sections S1.3 and S1.4.2).

**TABLE 1.**
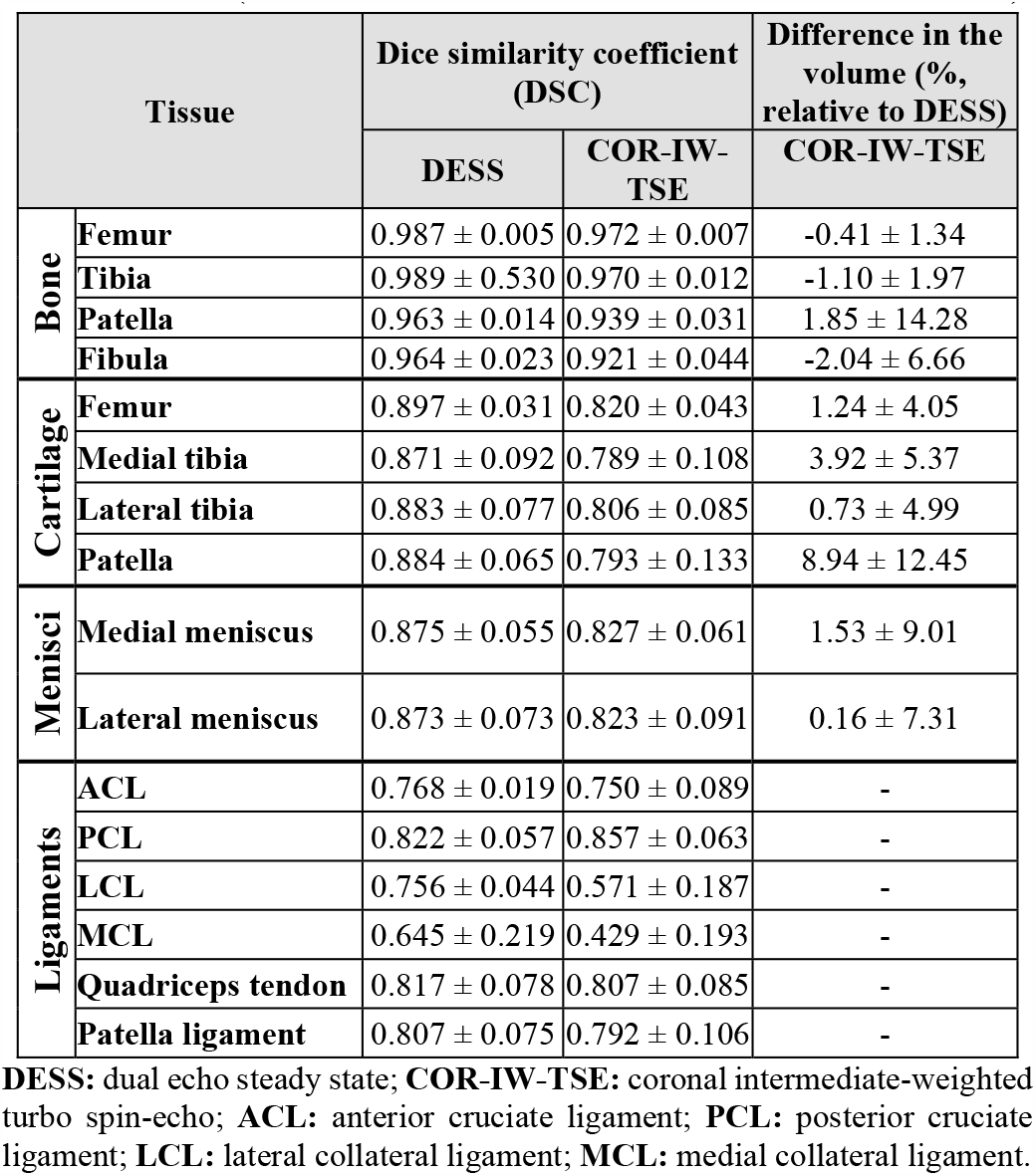
Dice similarity coefficient and the difference in tissue volumes (manually segmented vs. auto-segmentation). Magnitudes are mean ± standard deviation. No significant differences were observed between the tissue volume estimated by DESS and COR-IW-TSE images (*P*>0.05, independent-Samples Kruskal–Wallis test).

### G. Material assignment and model assembly

Within the MSK model, knee contacts were defined using a nonlinear elastic foundation model [37], [38] with Young’s modulus of 5 MPa and Poisson’s ratio of 0.45 for cartilage-cartilage and Young’s modulus of 3 MPa and Poisson’s ratio of 0.45 for meniscus-cartilage [30], [31].

The FRPVES material model was assigned to the cartilages and menisci within the template FE model (section II.D) using our in-house script, verified in our previous studies [6], [12], [17]. Notably, morphing the template did not require updates to the FRPVES material implementation, as morphing affected only nodal coordinates while preserving the mesh structure [17], [21]. The same script was also used to automatically assign the FRPVES material to the FE models generated by the auto-meshing algorithm (section II.E.1). Collagen fibril orientation and density and proteoglycan and fluid content were calculated for each element within the mesh and stored in ASCII format to be used in the Abaqus UMAT subroutine. Tissue composition and structure were obtained from the literature as detailed in the supplementary material.

The parts within the MSK and FE models were assembled to have ∼0.1 mm overclosure within the MSK model and ∼0.1 mm gap in the FE model. The overclosure in the MSK model was required for stable contact detection [38]. The tibiofemoral joint was assembled by fixing the femur and moving the menisci and tibia in the distal-proximal direction, while the patellofemoral joint was assembled by fixing the femur and moving the patella in the anteroposterior direction. The femur was fixed as the knee joint coordinate was defined with respect to the parent body (i.e., femur). We used the collision detection filter from the VTK Python library [51], and the parts were moved until reaching ∼0.1 mm overclosure or gap, correspondingly. Ligament insertion points were also translated according to their corresponding connected tissue.

### H. Simulations and statistical analysis

The segmentation accuracy was examined using the Dice Similarity Coefficient (DSC) for validation sets (Table 1) [69]. In addition, the output of the auto segmentation when using different sequences (i.e., DESS and COR-IW-TSE) for the same knee was evaluated by comparing the tissue volumes. In this analysis, DESS and COR-IW-TSE from all 507 MRIs from the ZIB dataset were segmented. Subsequently, volumes from each tissue were isolated by multiplying their total number of voxels by voxel volume. Last, we reported differences in tissue volumes between DESS and COR-IW-TSE (Table 1) and assessed if these differences were statistically significant. The comparison was made using the nonparametric Kruskal-Wallis analysis of variance test. The significance level was *p*<0.05.

To evaluate the cartilage thickness measurements, we included all seven KOA subjects and seven randomly selected healthy subjects from the fourth dataset of this study. We then compared the average thickness from the load-bearing regions of femoral and tibial cartilage, as well as from patellar and femoral groove cartilages (Fig. S3). Subsequent statistical analyses were the nonparametric Kruskal-Wallis analysis of variance, followed by a multiple-comparison corrected post-hoc test, to determine whether cartilage thickness differs between healthy and KOA subjects. The significance level was set *a priori* to *p*<0.05. All statistical tests were performed using MATLAB statistical toolbox (v2022b).

Motion analysis data were not available for the datasets of this study. Consequently, we ran a passive knee flexion test to evaluate the MSK models. We used 52 MSK models from the fourth dataset to estimate secondary knee kinematics and compared them against the literature [30]. Also, we performed the same analysis but using the MSK models assembled without menisci. In the literature, neglecting the menisci in MSK model assembly is common, as the inclusion of menisci substantially exacerbates convergence issues and computational costs.

In addition to the passive knee motion simulations listed above, tissue mechanics within the knee cartilage were estimated during walking gait using the FE models. Specifically, the loading and boundary conditions during stance phase of walking were applied to both template-based and auto-meshing FE models. The specifics of the loading and boundary conditions were obtained from the literature [9], and they were identical for all models. They consisted of knee flexion angle, tibiofemoral and patellofemoral joint contact forces, and knee varus-valgus and internal-external moments. Abaqus (v2022, Dassault Systèmes, US) soil consolidation solver was used. Due to time-consuming FE simulations (∼25 hours per model), we ran FE analysis for 15 randomly selected subjects. The KOA subjects were excluded from FE analysis, as the template-based model could not account for cartilage lesions (while the auto-meshing method could consider lesions). If the subject had a cartilage lesion (e.g., localized crack or defect), tissue mechanics could substantially differ between template-based and auto-meshing methods due to stress concentration around the lesion [70]. We presented the peak of maximum principal stress (i.e., maximum tensile stress), collagen fibril strain, and maximum shear strain within the cartilages. These quantities have been suggested to govern cartilage degradation, e.g., collagen network damage, proteoglycan loss, and cell death [6]–[8], [12]. The estimated tissue mechanics were compared point-by-point (as a function of time) using statistical parametric mapping (SPM) paired t-tests [71], with *p*<0.05.

The template-based [9], [17], [23], [72], [73] and auto-meshing [22], [29] algorithms have previously been compared to manual segmentation and meshing. In consideration of these prior studies and the significant time and effort involved in manual segmentation and modeling, we did not conduct a comprehensive comparison between manually created models and those of our study. Nonetheless, we manually created an FE model for a randomly selected healthy subject (from the fourth dataset) and compared the estimated tissue mechanics with the template-based and auto-meshing models of the same subject.

We tested the segmentation and modeling tool and conducted all the MSK and FE analyses on a workstation with an Intel Xeon Gold 6248 (involving eight threads per analysis) and an NVIDIA RTX 3060 (involving <5GB of vRAM memory).

## III. Results

It took ∼10 min to create one subject’s MSK and FE models (both template-based and auto-meshing) from the MRIs without user interaction needed. The process involved auto-segmentation (∼3 min), processing the geometries (∼2 min), and creating template-based (∼2 min) and auto-meshing (∼3 min)

MSK and FE models. Inputs were knee MRI in NIFTI format, and outputs were two sets of ready-to-use MSK and FRPVES FE models (template-based and auto-meshed). Out of 52 knee joints (26 left and 26 right), the template-based method had no failures, and the auto-meshing method had ∼85% success rate, with 8 failures mostly due to a few poorly generated elements in menisci leading to non-convex hull cross-sections.

Tissue volumes between auto-segmentation results obtained from DESS and COR-IW-TSE sequences were not significantly different (Table 1, datasets 1 to 3 of the study). When using PDW-VISTA-SPAIR images (DESS approximate) from the fourth study dataset, the average thickness of the load-bearing region from the medial femoral and tibial cartilages, as well as medial patellar and femoral groove cartilages, was significantly lower (*p*<0.05) in those with KOA (i.e., OA-affected knee) compared to healthy subjects (Fig. 2). Similar findings were also observed when comparing the contralateral (i.e., unaffected/less affected) knee of KOA subjects to healthy subjects, except for the medial tibia and femur (Fig. 2). The average thickness of load-bearing regions was not significantly different on the lateral femoral and tibial cartilages between KOA subjects and those of healthy subjects (Fig. 2).

From the passive knee flexion, secondary knee kinematics estimated by template-based and auto-meshing MSK models fell within the same range as literature data [30] (Figs. 3 and 4). The root mean square error (RMSE) between the secondary knee kinematics estimated by the template-based and auto-meshing MSK models, either with or without menisci, was <2.3° for tibiofemoral abduction/adduction rotation, <3.3° for internal/external rotation, <2.0 mm for anteroposterior translation, <2.8 mm for distal/proximal translation, and <0.9 mm for mediolateral translation (Figs. 3 and 4). The RMSE of the estimated secondary knee kinematics between MSK models with and without menisci, either generated by template-based or auto-meshing (Figs. 3 and 4), across the entire passive tibiofemoral flexion range was <0.7° for abduction/adduction rotation, <2.2° for internal/external rotation, <1.4 mm for anteroposterior translation, <0.8 mm for distal/proximal translation, and <0.7 mm for mediolateral translation.

**Fig. 3.**
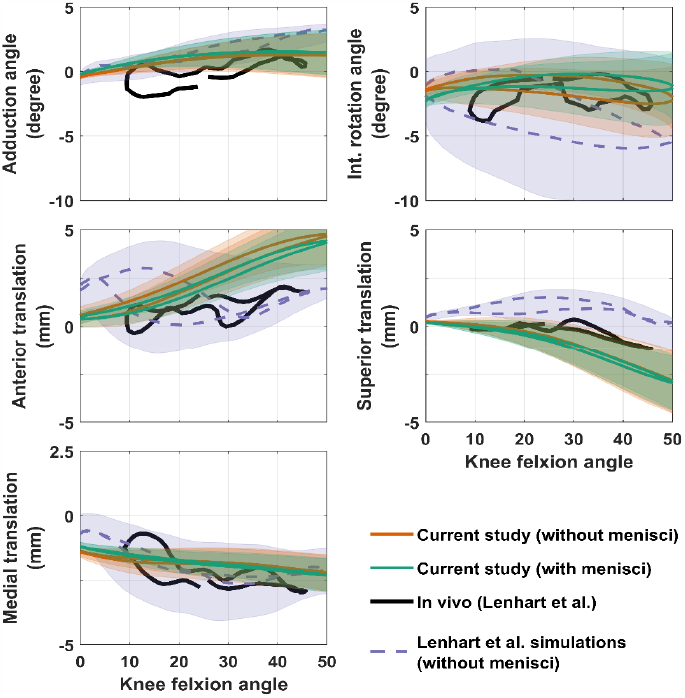
Secondary knee kinematics during passive knee flexion estimated by the template-based MSK models of our study without menisci (in orange) and with menisci (in green), compared to those from Lenhart et al. simulations [30] (in violet) and *in vivo* measurements [30] (in black). Highlights show within-subject variations, as well as variations due to uncertainties in ligament material parameters from Lenhart et al. [30].

**Fig. 4.**
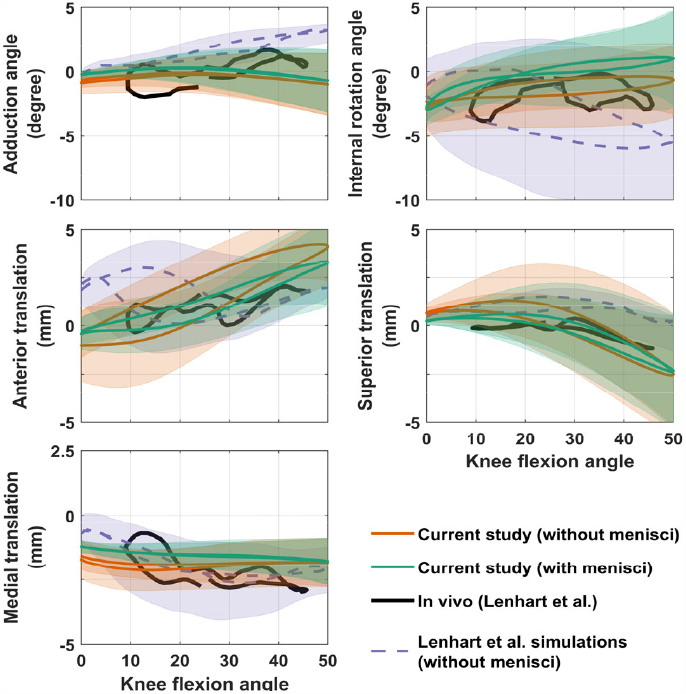
Secondary knee kinematics during passive knee flexion estimated by the auto-meshing MSK models of our study without menisci (in orange) and with menisci (in green), compared to those from Lenhart et al. simulations [30] (in purple) and *in vivo* measurements [30] (in black). Highlights show within-subject variations, as well as variations due to uncertainties in ligament material parameters from Lenhart et al. [30].

**Fig. 5.**
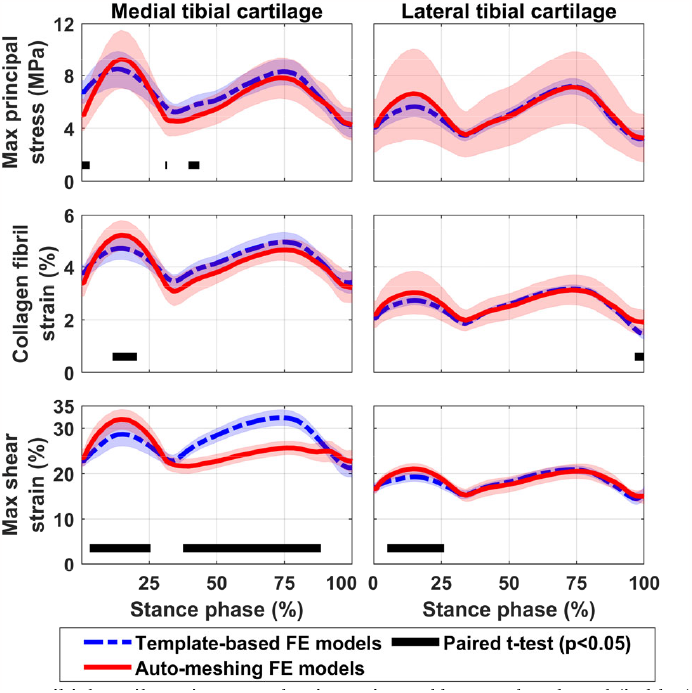
Tibial cartilage tissue mechanics estimated by template-based (in blue) and auto-meshing (in red) finite element (FE) models using PDW-VISTA-SPAIR (DESS approximate) images of fourth dataset of the study. Highlights show ±1 standard deviation, and black lines show time points at which results from the two FE modeling methods were statistically different

**Fig. 6.**
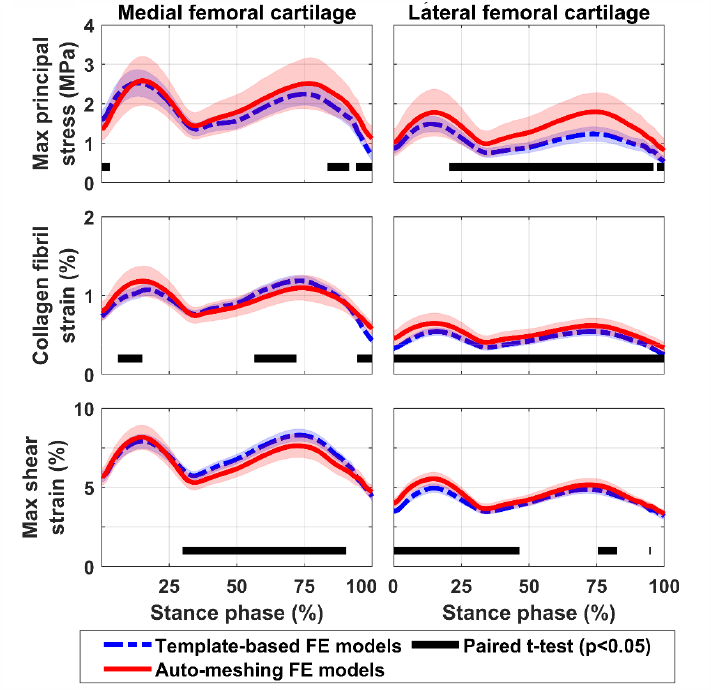
Femoral cartilage tissue mechanics estimated by template-based (in blue) and auto-meshing (in red) finite element (FE) models using PDW-VISTA-SPAIR (DESS approximate) images of fourth dataset of the study. Highlights show ±1 standard deviation, and black lines show time points at which results from the two FE modeling methods were statistically different.

Both the pattern and magnitude of tissue mechanics estimated by the template-based FE models agreed with those estimated by the auto-meshing FE models. Within the medial and lateral tibial cartilages (Fig. 5), RMSE between two FE modeling methods for maximum principal stress was 1.5 MPa (medial) and 2.3 MPa (lateral), for collagen fibril strain 0.6 percentage points (medial) and 0.53 percentage points (lateral), and for maximum shear strain 5.1 percentage points (medial) and 1.75 percentage points (lateral). Within the medial and lateral femoral cartilages (Fig. 6), RMSE between two FE modeling methods for maximum principal stress was 0.4 MPa (medial) and 0.5 MPa (lateral), for collagen fibril strain 0.12 percentage points (medial) and 0.11 percentage points (lateral), and for maximum shear strain 0.76 percentage points (medial) and 0.33 percentage points (lateral). Within medial and lateral patellar cartilages (Fig. 7), RMSE of tissue mechanics between methods for maximum principal stress was 0.86 MPa (medial) and 0.80 MPa (lateral), for collagen fibril strain 0.11 percentage points (medial) and 0.06 percentage points (lateral), and for maximum shear strain 0.65 percentage points (medial) and 0.45 percentage points (lateral). The estimated tissue mechanics, both magnitude and distribution, were comparable between the manually generated, template-based, and auto-meshing FE models of the randomly selected subject (Fig. 8, maximum principal stress at the knee contact force peak during walking).

**Fig. 7.**
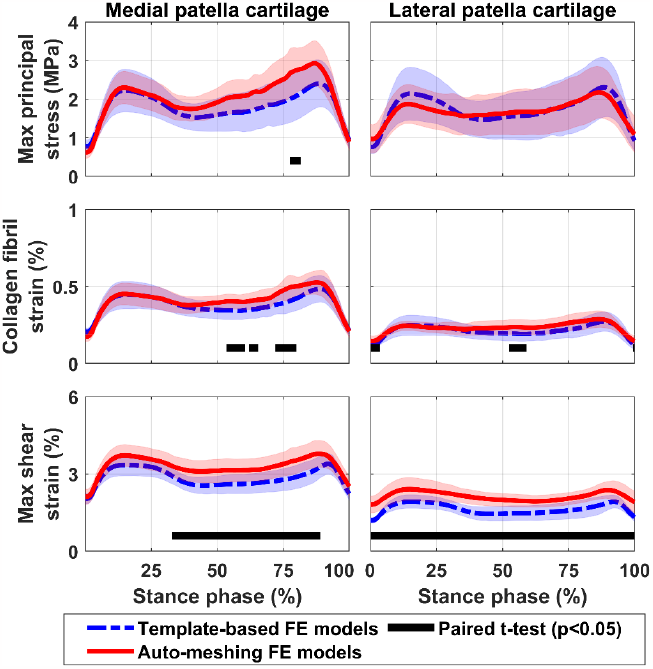
Patellar cartilage tissue mechanics estimated by template-based (in blue) and auto-meshing (in red) finite element (FE) models using PDW-VISTA-SPAIR (DESS approximate) images of fourth dataset of the study. Highlights show ±1 standard deviation, and black lines show time points at which results from the two FE modeling methods were statistically different.

**Fig. 8.**
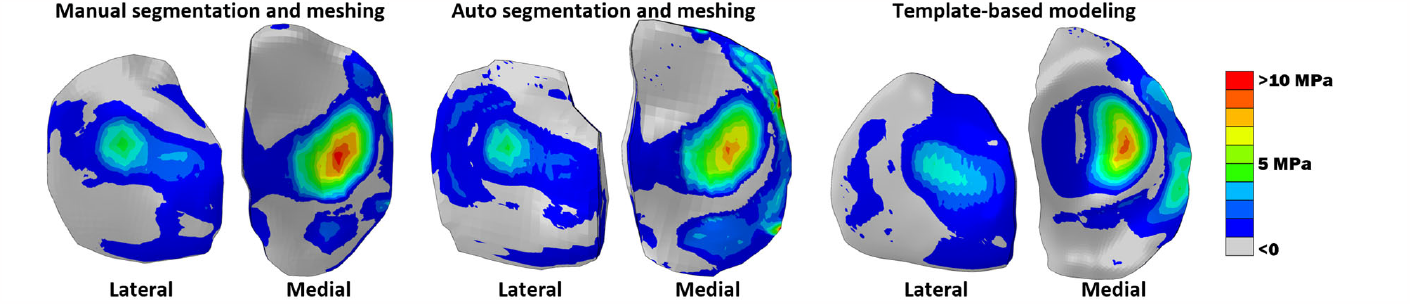
Maximum principal stress within tibial cartilage at the peak of knee joint contact force during walking, estimated by the manually created (left), auto-segmented and meshed (middle), and template-based (right) finite element models of a randomly selected subject.

## IV. Discussion

### A. Summary

This study is the first to develop a fully automated and robust knee MSK and FE modeling tool. The tool requires no interactions from the user and, in <10 min, performs preprocessing of MRI, auto segmentation of bones, cartilages, menisci, and the major knee ligaments, and creates two sets of knee MSK and FE models, i.e., template-based and auto-meshing. Moreover, the tool provided consistent integration between MSK and FE models of the knee [14], which has previously demonstrated great potential for predicting person- and task-specific knee kinematics [9], [17], [74], kinetics [9], [14], [17], [74], and tissue-level mechanics [6], [9], [14], [17]. The MSK model incorporated personalized knee ligaments and contact geometries, including menisci, with a nonlinear elastic foundation contact. The FE model incorporated a detailed FRPVES material model, previously used to predict mechanobiological degradation of knee cartilage, e.g., collagen damage, proteoglycan loss, and cell death [6], [7], [12]. Overall, we observed minor discrepancies in knee biomechanics between the models created from different MRI sequences as well as the template-based and auto-meshing approaches.

### B. Image segmentation

The DSC for bones, cartilages, and menisci obtained from auto-segmentation were of the same range as those reported in the literature [22], [75]–[78]. Notably, 3D nnU-Net showed comparable DSC but fewer segmentation artifacts than 2D nnU-Net. Kessler et al. [79] also reported higher accuracy in extracting cartilage morphology and composition when using 3D compared to 2D U-Nets. Moreover, the tool developed in our study demonstrated the potential to segment different MRI sequences and datasets, which is critical for tool applicability especially for clinical use. We observed lower DSC for COR-IW-TSE compared to DESS sequence (Table 1), which can be attributed to the noticeably lower image resolution in the COR-IW-TSE (see section II.A and Fig. S2). Although auto-segmentation from COR-IW-TSE sequence slightly overestimated tissue volumes (Table 1) and thicknesses (Figs. 2 vs. S5) compared to auto-segmentation from DESS, segmented tissue volumes were not significantly different (p>0.05) between sequences (Table 1). Also, auto-segmentation using COR-IW-TSE sequence better segmented knee ligaments (including LCL and MCL) compared to DESS, potentially due to higher contrasts between ligaments and surrounding tissue in COR-IW-TSE compared to DESS sequences (also addressed by Peterfy et al. [41]).

We tested the modeling tool with a dataset consisting of healthy and KOA subjects. The cartilage thickness obtained from the auto-segmentation of the OA-affected tissue was significantly lower than that of healthy subjects (Fig. 2). This was in line with the medical history of the patients, as KOA subjects had a previously diagnosed joint space narrowing. Also, the patellar cartilage thickness, as well as femoral groove cartilage thickness, were significantly lower in the KOA compared to healthy subjects (Fig. 2). This is understandable based on the fact that medial tibiofemoral OA is often combined with patellofemoral OA [80].

### C. MSK and FE modeling

Considering the range of knee kinematics (Figs. 3 and 4), the secondary knee kinematics of passive knee flexion estimated by the MSK models, with and without menisci, were in good agreement with *in vivo* measurements from literature as well as simulations by Lenhart et al. [30] (Figs. 3 and 4). Notably, the MSK models of our study were generated based on the models developed by Lenhart et al. (menisci excluded) [30] and Smith et al. (menisci included) [31]. Both these MSK models have been developed and adjusted to produce secondary knee kinematics and knee contact forces comparable with *in vivo* data [30], [31]. Nevertheless, including the menisci within the MSK model added 12 DoF to the knee joint, resulting in a total of 24 DoF joint. This significantly increased the computational costs and convergence issues. Our results, in line with those of Lenhart et al. [30], suggest that if menisci contact mechanics (e.g., contact pressures) are not of interest, menisci might be excluded from the MSK analysis as they do not influence knee kinematics but substantially add to computational burdens.

Overall, the estimated tissue mechanics within medial and lateral tibial, femoral, and patellar cartilages had a consistent pattern, magnitude, and within-subjects variations (shaded area in Figs. 5 to 7 and S6 to S8) between the auto-meshing and template-based FE models. In certain instances, the auto-meshed FE models exhibited greater within-subject variations in comparison to the template-based FE models. Despite receiving identical inputs, the distinct within-subject variations observed can be attributed to the differing capabilities of the template-based and auto-meshing approaches to represent the knee geometries of the subjects. Also, we observed a few cases with noticeable differences in the magnitude, e.g., maximum shear strain in the medial tibial cartilage (Fig. 5). However, both the auto-meshing and template-based FE modeling approaches estimated comparable peak values during the stance phase.

The deviation in the peak of the estimated tissue mechanics between template-based and auto-meshing FE models (magnitudes) was <1.5 MPa for maximum principal stress, <1 percentage point for collagen fibril strain, and <3 percentage points for maximum shear strain. Also, differences in the magnitudes of knee cartilage mechanics between the FE models from COR-IW-TSE and DESS images, either auto-meshing or template-based FE models, were marginal, i.e., <0.6 MPa for maximum principal stress, <0.8 percentage points for collagen fibril strain, and <2 percentage points for maximum shear strain (Figs. 5 to 7 vs. S6 to S8). Although these differences occasionally reached statistical significance during the gait stance (*p*<0.05, solid black lines in Figs. 5-7 and S6-S8), they were considerably lower than the cartilage degradation thresholds, e.g., 7 to 30 MPa for tensile stress [81] and ∼30% to 50% for maximum shear strain [6], [12]. This implies a consistent prediction of cartilage degradation using either template-based or auto-meshing pipelines from the MRI sequences tested in the current study. Our previous studies have also demonstrated comparable cartilage mechanics and degradation between template-based and manual FE modeling [21], [82]. Results of the current study suggest that the template-based method is viable, particularly because it showed no failures to create MSK and FE models and had substantially fewer convergence issues than the auto-meshing method.

The template-based [9], [17], [23], [72], [73] and auto-meshing [22], [29] algorithms adopted in our study have been previously investigated against manual modeling and found to provide consistent tissue volume and tissue mechanics [22], [29]. Here, we manually created an FE model from a randomly selected healthy subject and observed comparable tissue mechanics with auto-meshing and template-based models (Fig. 8).

### D. Limitations

The number of knees with segmented ligaments in this study was limited (i.e., dataset three). To mitigate this effect on the training of auto-segmentation neural networks, we applied rich data augmentation techniques. Additionally, this dataset was only used for ligament insertion point extraction and no other segmentations. In support of ongoing research, we are currently performing additional manual segmentations of the knee [83] to improve the auto-segmentation of knee ligaments. In this study, we did not focus on cartilage and menisci defects such as cracks or total loss, as we developed the modeling tools first to focus on subjects without or at early-stage KOA. However, the auto-meshing algorithm used in our study has shown the potential to consider cartilage lesions [22], and the template-based method employed an FFD algorithm with the potential to account for cartilage thickening/thinning typically observed in early KOA. Despite the promising features, the auto-meshing algorithm in its current form needs further modifications and adjustments to ensure a valid mesh around the localized osteochondral lesions. Muscle paths and parameters (e.g., strength) within the MSK model were not personalized, as this would require additional data (unavailable in our study datasets) and extensive validation beyond the scope of this study. Our future studies aim to update the insertions and wrapping surfaces of muscles spanning the knee using the segmented knee bones. We also plan to personalize muscle strength according to muscle volumes from MRI or, alternatively, through optimization while subjects perform different exercises at known exertions (e.g., maximal contractions). Also, the models currently lack personalized material parameters for cartilages, menisci, and ligaments. To the authors’ knowledge, no methods are available to extract personalized material parameters (especially for the FRPVES material model) for the whole knee tissue non-invasively. Our future studies aim to automatically extract and assign subject-specific ligament [84] and cartilage material parameters [85].

### E. Applications and future developments

Several powerful modeling tools have been recently published to support the study of knee biomechanics [17], [21]– [23]. However, these tools are not fully automated [17], [21], [23], do not include whole tibiofemoral and patellofemoral joints (such as menisci) [21], [22], or do not offer MSK modeling [21]–[23]. The tool developed in this study, which has integrated MSK and FE models, can be applied to the study of knee biomechanics during different functional activities and rehabilitation exercises or knee injuries/surgeries such as ligaments and menisci [9], [31], [86]. More specifically, the FE model with FRPVES material incorporates the structure and composition of knee cartilage and menisci (such as collagen network and proteoglycan and water content) and their interplay with tissue mechanobiological response governing tissue degradation and remodeling [2], [3].

The developed tool is modular to facilitate its customization. For example, it is readily re-deployed with different MRI segmentation algorithms, geometric processing, meshing, etc. Moreover, the algorithms developed in this study, e.g., the template-based method, can be adapted for automated modeling of other joints, such as the hip, ankle, shoulder, and vertebral column. Importantly, the template-based method could generate MSK and FE models even without auto-segmentation data [23]; however, we used auto-segmented cartilages and menisci as regions of interest to enhance image registration (section II.D). The cartilage thickness map also has potential for further research and clinical applications (e.g., computer-aided diagnoses). Our future studies will use the cartilage thickness map to extract the cartilage strain map and personalize cartilage material parameters or validate the MSK and FE models. Moreover, our future research extensively focuses on integrating the tool developed in this study with out-of-lab motion analysis [87] and mechanobiological cartilage degradation algorithms [6], [12]. We aim to introduce a clinically viable tool to predict the onset and progression of KOA and provide personalized rehabilitation and gait modification to optimize knee biomechanics in the interest of preventing or decelerating KOA progression [6], [9], [12].

## V. Conclusion

We introduced a novel, fully automated, easy-to-use, and robust knee MSK and FE modeling tool with a GUI, which enables personalized and rapid (several minutes) knee modeling for research or clinical applications with large cohorts. The developed tool demonstrated the potential to accurately extract knee geometries from different MRI sequences. Importantly, the MSK and FE models generated by the automated tool estimated knee biomechanics consistent with those reported in the literature. Our future studies will integrate the auto segmentation and MSK-FE modeling tool with a cartilage degradation algorithm to predict task- and person-specific knee cartilage degradation response, enabling personalized rehabilitation and gait modification to delay KOA progression.

## Appendix

More information on the study is provided in the supplementary material. The developed tool is available from the corresponding author upon request.

## Notes

This study was financially supported by the National Health and Medical Research Council (NHMRC) Idea Grant (Award No: APP2001734), the Academy of Finland (Grant No: 322424, 324529), Sigrid Juselius foundation.

### Competing Interest Statement

The authors have declared no competing interest.

