## Supplementary Materials for "An Automated and Robust Tool for Musculoskeletal and Finite Element Modeling of the Knee Joint"

### S1. Methods

#### S1.1. Knee coordinate system

The knee joint anatomical coordinate system (ACS) was used from the template-based approach, as we found this method more robust than fitting-based approaches, e.g., by Miranda et al. [1]. Here, we first determined the knee ACS of the template knee using the method introduced by Miranda et al. [1]. Next, we obtained the transformation matrix of the template model, which translates the posterior axis of the ACS to +X, the proximal axis of the ACS to +Y, and the mediolateral axis of the ACS to Z (Fig. 1-C, right-handed coordinate systems). This transformation matrix was then used to translate the subjects' geometries from template-based as well as auto-segmentation methods to the knee coordinate system. NiftyReg tool [2] was used to obtain transformation matrices, as explained in section 2.4 of the manuscript.

Nevertheless, the knee flexion angle within the magnetic resonance images (MRIs) often varies across the subjects. To consider this, we performed two affine registrations. We first registered the subject's MRI to the template MRI using tibial cartilage and menisci as the mask. The registered image was then re-registered to the template MRI but using only the femoral cartilage as the mask. The rotation around the flexion-extension axis between the two transformation matrices was considered the difference between the template's and the subject's knee flexion angles and was applied to the subject's MSK-FE models (i.e., to the tibia, patella, menisci, and the corresponding ligament insertion points). Notably, the MRI from the template subject (i.e., the template MRI) was taken with the knee at full extension.

The knee joint in both the MSK and FE models had identical geometries and coordinate systems. The MSK model had a knee with tibiofemoral, patellofemoral, and medial and lateral meniscus joints [3]. Each joint had 6 degree-of-freedom (DoF), with a coordinate system identical (i.e., at full extension) to the knee

joint ACS, explained above. In the MSK models, all the coordinate systems were fixed relative to the tibia except for the patella, which was fixed relative to the femur [3]–[5]. The MSK model without the menisci had no medial and lateral meniscus coordinates.

A global coordinate system was defined within the FE models, identical to the knee ACS. Then we defined a local coordinate system between the patella and femur, consistent with the patellofemoral joint within the MSK model. Menisci had no separate (i.e., local) coordinate systems, as they were free to move and were governed by the contact between the femoral and tibial cartilages and menisci horn attachments. All the DoF of nodes on the tibial cartilage-subchondral bone interface were closed, and the inputs to the FE models were applied to the femur (relative to the tibia) and patella (relative to the femur), consistent with the MSK models. The FE model coordinate system, loading, and boundary conditions have been extensively explained in our previous studies [6], [7].

#### S1.2. Cartilage thickness mapdid

We extracted the femoral, tibial, and patellar cartilage thickness maps (Figs. 1-B and S3). Using the Visualization Toolkit Python library (VTK v9.2) [8], a ray was cast from each vertex on the bones in the direction of their respective normals. If the ray had two intersections with the corresponding cartilage geometry (within an arbitrarily-chosen ±1 cm range), the distance between the two intersections was considered cartilage thickness, and the value was assigned to that vertex on the bone. Consequently, the cartilage thickness map had a resolution equal to the number of bone vertices (Figs. 1-B and S3). Lastly, we calculated the average cartilage thickness for the load-bearing regions of medial and lateral femoral, tibial, and patellar cartilages (Fig. S3). Cartilage thickness maps were extracted before applying the Taubin and Laplacian smoothing filters (see section 2.3 of the manuscript).

It was a challenge to detect regions with total cartilage loss from the surrounding bone, as both regions have zero cartilage thickness (e.g., blue regions in Fig. S3). To overcome this challenge, first, we obtained an initial estimate for the geometry of the subjects' cartilages, i.e., healthy cartilages, before any loss occurred. The scaled template model was used as this initial estimate (see section 2.4 of the manuscript). Next, the subjects' cartilage (Fig. S3), including regions with total cartilage loss, was considered as the sum of the region obtained from the scaled template (thickness set to zero) and the region obtained from auto-segmentation (thickness >0).

Finally, the cartilage thickness was extracted from the load-bearing regions of the medial and lateral femoral and tibial cartilages, femoral groove, and medial and lateral patella cartilages. The regions' definition was adopted from our previous study [9], with details on obtaining each region within the knee cartilage demonstrated in Fig. S3. Lastly, the average cartilage thickness (reported in Figs. 2 and S6) was obtained for the load-bearing regions of the medial and lateral femoral and tibial cartilages, femoral groove, and medial and lateral patellar cartilages.

#### S1.3. Insertion points of ligaments, quadriceps tendon, and menisci horn attachments

All the MSK-FE models, either template-based or auto-meshing approach, had identical ligament insertion points, stiffness, and slack length. The insertion points of the anterior cruciate ligament (ACL) and posterior cruciate ligament (PCL) on the femoral and tibial bones, as well as the insertion points of the patella ligament and quadriceps tendon on the tibial and patella bones, were determined using the nearest-neighbor concept. In this, the points on the femur, tibia, and patella bones with less than 2 mm distance from each end of the aforementioned ligaments (from auto-segmentation) were considered insertion points, corresponding to distal and proximal ends.

The auto-segmentation of lateral and medial collateral ligaments (LCL and MCL) was relatively poor (Table 1); thus, the insertion points of LCL, MCL, and lateral and medial patellofemoral ligaments (LPFL, and MPFL) were obtained from the segmented bones, according to the literature (Fig. 1-D) [4], [10]–[14]. For LCL, MCL, and femoral insertions of LPFL and MPFL, we selected the insertion point region from the corresponding bone geometry (i.e., femur, tibia, or fibula) according to the cross-section area and insertion point from the literature [4], [10]–[14]. To obtain the patella end of LPFL and MPFL, first, two vertices on the very end of the patella bone in medial and lateral directions were found. Then, two rectangular surfaces, centered at the selected vertices with dimensions from [10], were selected from the patella bone and considered LPFL and MPFL insertion region (Fig. 1-D).

Then, each chosen region of the bone meshes was re-meshed to have 30 vertices for ACL, PCL, MCL, and LCL, and 6 vertices for LPFL, MPFL, and patella ligament. Lastly, spring elements were defined between the vertices for each of the ligaments.

If the algorithm failed to detect the insertion of some ligaments, the corresponding insertions were used from the scaled template model. We used this approach to increase the robustness of the modeling, especially when the segmentation of the ligaments was suboptimal. Also, the menisci horn attachments on the tibia bone were used from the scaled template model. The insertions of the remaining ligaments, e.g., popliteofibular ligament and posteriomedial capsule within the MSK model, were untouched. The slack length and stiffness of all the ligaments were set according to Lenhart et al. [5] and Smith et al. [3], and menisci horn attachments according to Villegas et al. [15].

### S1.4. The material models of cartilages, menisci, and ligaments

#### S1.4.1. Cartilage and menisci

A fibril-reinforced poroviscoelastic swelling (FRPVES) material model [16], [17] was utilized for the cartilages, and the depth-dependent Benninghoff-type (arcade) architecture of collagen fibers was implemented as split-lines for femoral, tibial, and patellar cartilages [18]–[21]. Menisci were modeled as a fibril-reinforced poroelastic swelling (FRPES) material.

Specifically, the FRP(V)ES model assumes that the tissue is composed of solid and fluid matrices. The solid matrix is separated into a porous nonfibrillar part, representing the proteoglycan matrix, and a fibrillar network (viscoelastic in cartilage and elastic in meniscus), describing the collagen fibrils, and the influence of swelling caused by fixed charge density (FCD) of proteoglycans. The total stress in the FRP(V)ES model ($\boldsymbol{\sigma}_{\mathrm{t}}$) is given by:

$$\boldsymbol{\sigma}_{\mathrm{t}} = \boldsymbol{\sigma}_{\mathrm{nf}} + \boldsymbol{\sigma}_{\mathrm{f}} - \Delta\pi\mathbf{I} - \mu^{\mathrm{f}}\mathbf{I} - \mathrm{T_c}\mathbf{I} \quad \text{(Eq. S1)}$$

where $\boldsymbol{\sigma}_{\mathrm{nf}}$ and $\boldsymbol{\sigma}_{\mathrm{f}}$ are the stress tensors of the nonfibrillar and fibrillar matrices, $\Delta\pi$ is the swelling pressures, $\mathbf{I}$ is the unit tensor, $\mu^{\mathrm{f}}$ is the chemical potential of water, and $\mathrm{T_c}$ is the chemical expansion stress. The nonfibrillar matrix was modeled by compressible neo-Hookean properties. The stress within the nonfibrillar matrix is given by [22]:

$$\boldsymbol{\sigma}_{\mathbf{nf}} = \mathrm{K}\frac{\ln(\mathrm{J})}{\mathrm{J}}\mathbf{I} + \frac{\mathrm{G}}{\mathrm{J}}(\mathbf{F}\mathbf{F}^{\mathbf{T}} - \mathrm{J}^{2/3}\mathbf{I}) \quad \text{(Eq. S2)}$$

$$\mathrm{G} = \frac{\mathrm{E_m}}{2(1+\nu_{\mathrm{m}})} \quad \text{(Eq. S3)}$$

$$\mathrm{K} = \frac{\mathrm{E_m}}{3(1-2\nu_{\mathrm{m}})} \quad \text{(Eq. S4)}$$

where G and K are the shear and bulk moduli of the nonfibrillar matrix, J is the determinant of the deformation tensor **F**, $E_m$ and $\nu_m$ are Young's modulus and Poisson's ratio of the nonfibrillar matrix, respectively.

The fluid flow in the nonfibrillar matrix is simulated according to Darcy's law given by:

$$q = -k\nabla p \tag{Eq. S5}$$

where q is the flux in the nonfibrillar matrix, $\nabla p$ is the hydrostatic pressure gradient vector across the region, and k is the hydraulic permeability. The hydraulic permeability is defined to be strain-dependent [23], given by [24]:

$$k = k_0 \left(\frac{1+e}{1+e_0}\right)^M \tag{Eq. S6}$$

where $k_0$ is the initial permeability, e and $e_0$ are the current and the initial void ratio, and M is a positive constant.

The cartilage collagen fibers were modeled as a viscoelastic material. In the material model, a nonlinear spring (with the strain-dependent modulus $E_\varepsilon \varepsilon_f$) is in series with a linear dashpot (with the damping coefficient $\eta$). This nonlinear spring-dashpot system is in parallel with a linear spring (with the initial modulus $E_0$). Fibrils were assumed to only resist tension; thus, the collagen fibril stress of the cartilage was formulated as [25]:

$$\sigma_f = \begin{cases} -\frac{\eta}{2\sqrt{(\sigma_f - E_0\varepsilon_f)E_\varepsilon}}\dot{\sigma}_f + E_0\varepsilon_f + \left(\eta + \frac{\eta E_0}{2\sqrt{(\sigma_f - E_0\varepsilon_f)E_\varepsilon}} + \right)\dot{\varepsilon}_f & \text{for} \quad \varepsilon_f > 0 \\ 0 & \text{for} \quad \varepsilon_f \leq 0 \end{cases} \tag{Eq. S7}$$

where $\sigma_f$ and $\varepsilon_f$ are the fibril stress and strain, and $\dot{\sigma}_f$ and $\dot{\varepsilon}_f$ are the fibril stress and strain rates.

The collagen fibers within the menisci were modeled as linear elastic material. The menisci collagen fiber stress was formulated as [26]:

$$\sigma_f = \begin{cases} E_\varepsilon \varepsilon_f & \text{for} \quad \varepsilon_f > 0 \\ 0 & \text{for} \quad \varepsilon_f \leq 0 \end{cases} \tag{Eq. S8}$$

The collagen fiber network consisted of primary and secondary fibrils [27]. The primary collagen fibrils form a depth-dependent arcade-like structure [28], while the secondary fibrils are randomly organized in 13 different random orientations [27]. Secondary fibrils mainly replicate the inter-fibril connections and cross-links in the collagen network. Consequently, defining C as the amount of the primary fibrils with respect to the secondary fibrils and $\rho_z$ as the relative collagen density, the stresses are given by [27]:

$$\sigma_{f,p} = \rho_z C \sigma_f \tag{Eq. S9.a}$$

$$\sigma_{f,s} = \rho_z \sigma_f \tag{Eq. S9.b}$$

The Donnan osmotic swelling pressure at equilibrium is given by:

$$\nabla\pi = \phi_{int} RT \left( \sqrt{c_{FCD}^2 + 4\frac{(\gamma_{ext}^{\pm})^2}{(\gamma_{int}^{\pm})^2} c_{ext}^2} \right) - 2\phi_{ext} RT c_{ext} \tag{Eq. S10}$$

where $c_{FCD}$ is the fixed charge density content at equilibrium, $c_{ext}$ is the external salt concentration (0.15 M), $\phi_{int}$, $\phi_{ext}$, $\gamma_{int}^{\pm}$, and $\gamma_{ext}^{\pm}$ are internal and external osmotic coefficients and internal and external activity coefficients, respectively, R is the molar gas constant (8.3145 J/mol·K), and T is the absolute temperature (293 K). When temperature and external salt concentration are constant, then the only variable is the FCD which can be defined as a function of the tissue deformation, as:

$$c_{FCD} = c_{FCD0}\left(\frac{n_{fl0}}{f_{fl0}-1+J}\right) \quad \text{(Eq. S11)}$$

where $c_{FCD0}$ is the initial fixed charge density and $n_{fl0}$ is the initial fluid volume fraction. The chemical expansion stress $T_c$ is determined as:

$$T_c = a_0 c_{FCD} \exp\left(-\kappa \frac{\gamma_{ext}^{\pm}}{\gamma_{int}^{\pm}} \sqrt{c^-(c^- + c_{FCD})}\right) \quad \text{(Eq. S12)}$$

where $a_0$ and $\kappa$ are material constants [16] and $c^-$ is the mobile anion concentration.

**Table S1.** Material parameters for the knee joint cartilages and menisci [25], [27], [29]–[32]

| Material parameter | Description | Femoral cartilage | Tibial cartilage | Patellar cartilage | Menisci |
|---|---|---|---|---|---|
| $E_m$ (MPa) | Young's modulus of the nonfibrillar matrix | 0.215 | 0.106 | 0.505 | 0.500 |
| $\nu_m$ (-) | Poisson's ratio of the nonfibrillar matrix | 0.15 | 0.15 | 0.15 | 0.36 |
| $k_0$ ($\times 10^{-15}$ $m^4 N^{-1} s^{-1}$) | Initial permeability | 6 | 18 | 1.9 | 1.25 |
| M (-) | Exponential term of strain-dependent permeability | 5.09 | 15.64 | 15.93 | 5.09 |
| $\eta$ (MPa · s) | Viscosity of collagen fibrils | 1062 | 1062 | 1062 | - |
| $E_0$ (MPa) | Initial fibril network modulus | 0.92 | 0.18 | 1.88 | - |
| $E_f$ (MPa) | Fibril network modulus | - | - | - | 28 |
| $E_\varepsilon$ (MPa) | Strain-dependent network modulus | 150 | 23.6 | 597 | - |
| C (-) | The ratio of primary collagen fibers to secondary collagen fibers | 12.16 | 12.16 | 12.16 | 12.16 |
| $n_{f,eq}$ (mEq/ml) | Fluid fraction | *$0.85 - 0.15h$ | | | 0.72 |
| $c_{FCD0}$ (mEq/mL) | Initial depth-wise fixed charge density distribution | *$-0.1h^2 + 0.24h + 0.056$ | | | 0.03 |
| $\rho_z$ | Depth-wise collagen distribution | *$-20.6h^6 + 64.4h^5 + 78.1h^4 - 45.9h^3 + 13.4h^2 - 1.6h + 0.96$ | | | 1 |

*h: normalized cartilage depth from the surface

#### S1.4.2. Material model of ligaments and menisci horn attachments

Ligaments, quadriceps tendon, and menisci horn attachments were modelled as spring bundles to have sufficient accuracy in the estimated parameters while keeping the computational demand reasonable [33]. Utilizing a bundle of springs provides the ligament model with compression-tension nonlinearity with different properties in horizontal (along the length) and vertical (perpendicular to the fibrils/springs) directions.

All the ligaments and the quadriceps tendon within MSK-FE models were defined as nonlinear springs with no compressive resistance. The slack, toe, and linear regions of the ligament/tendon were formulated according to the study by Blankevoort et al. [34]:

$$
f = \begin{cases} 0 & \varepsilon < 0 \\ \frac{1}{4} K\ \varepsilon^2/\varepsilon_l & 0 \le \varepsilon \le 2\varepsilon_l \\ K\,(\varepsilon - \varepsilon_l) & \varepsilon > 2\varepsilon_l \end{cases} \qquad \text{(Eq. S13)}
$$

where f is the tensile force in each ligament/tendon element, K is the ligament stiffness, $\varepsilon_l$ represents the end of the toe region and was set to 0.03 [35], and ε is the strain in the ligament/tendon. All the ligament stiffnesses and slack lengths were selected according to those of Lenhart et al. [5]. The patella ligament had a stiffness of 545 N/mm [36]. Menisci horn attachments were modeled as linear spring bundles with a total stiffness of 336 N/mm and 381 N/mm for the anterior and posterior sides, respectively [15].

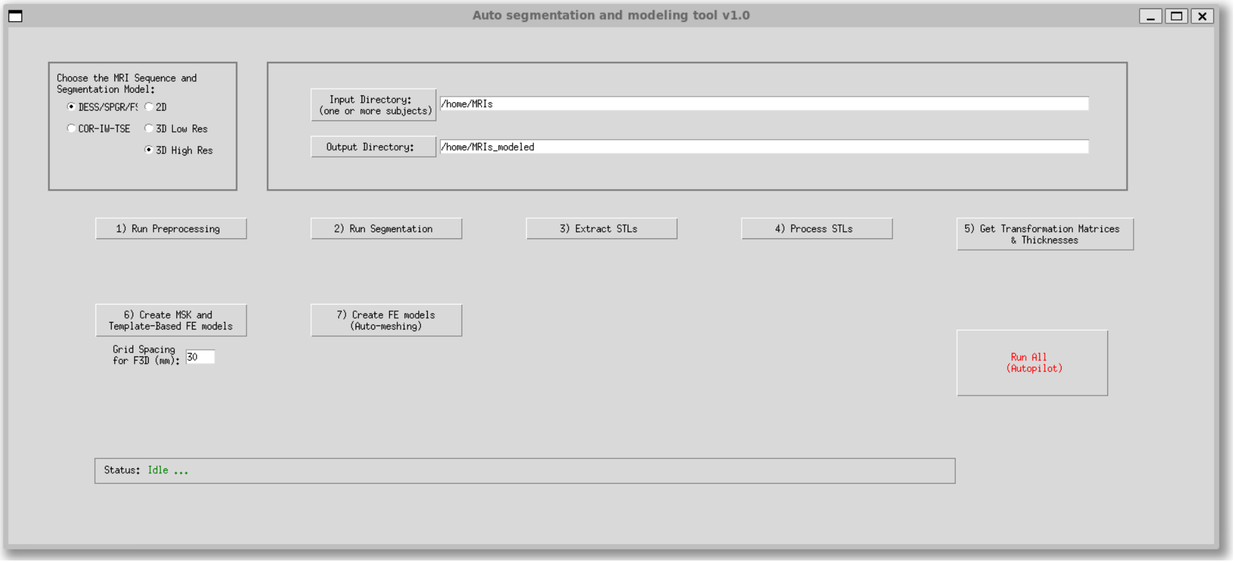


**Fig. S1.** The graphical user interface (GUI) of the developed tool.

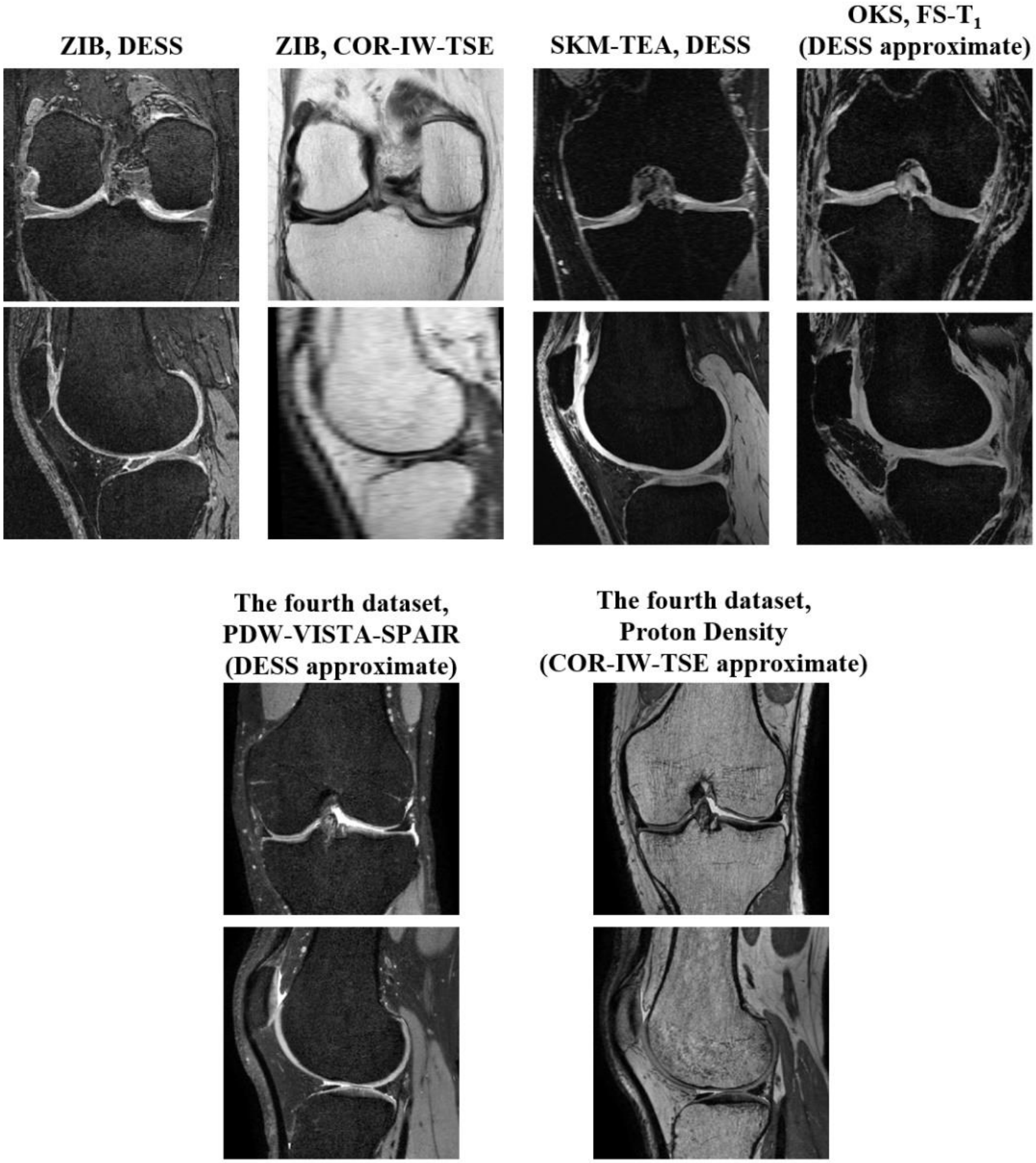


**Fig. S2.** Coronal and sagittal image slices from MRI sequences of the datasets used in the study (Top row: training dataset, Bottom row: testing dataset).

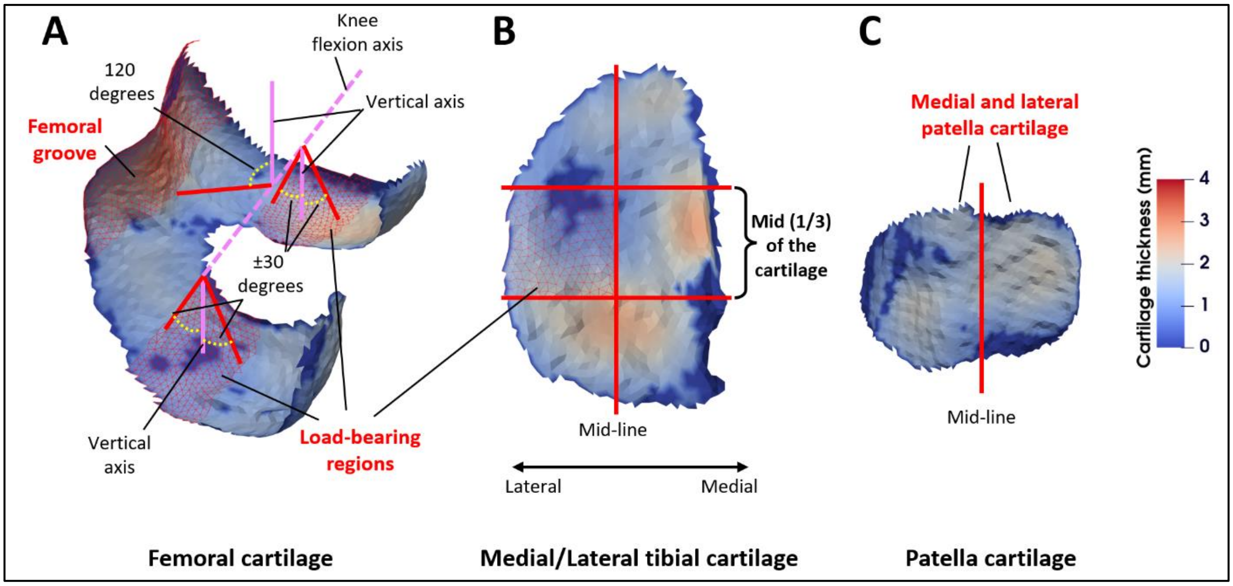


**Fig. S3.** Regional cartilage thickness for A) femoral cartilage, B) medial/lateral tibial cartilage, and C) patella cartilage.

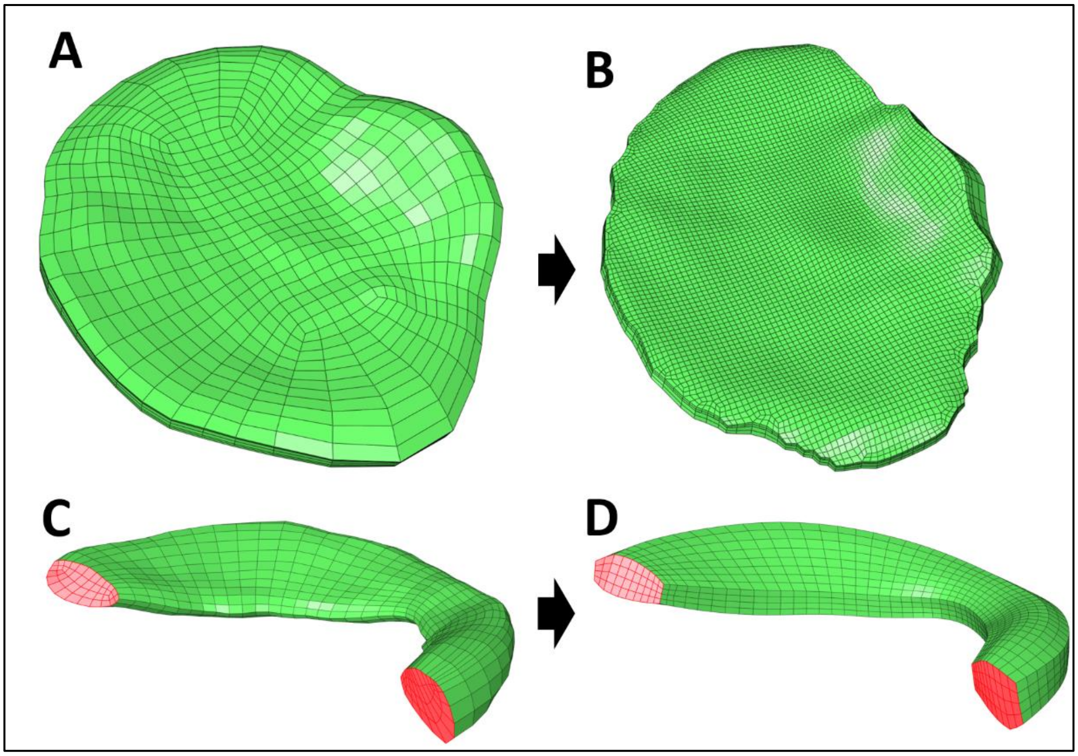


**Fig. S4.** The finite element mesh obtained from the algorithm developed by Rodriguez-Vila et al. [37] (A: Tibial cartilage, C: Meniscus) and the modified algorithm of our study (B: Tibial cartilage, D: Meniscus).

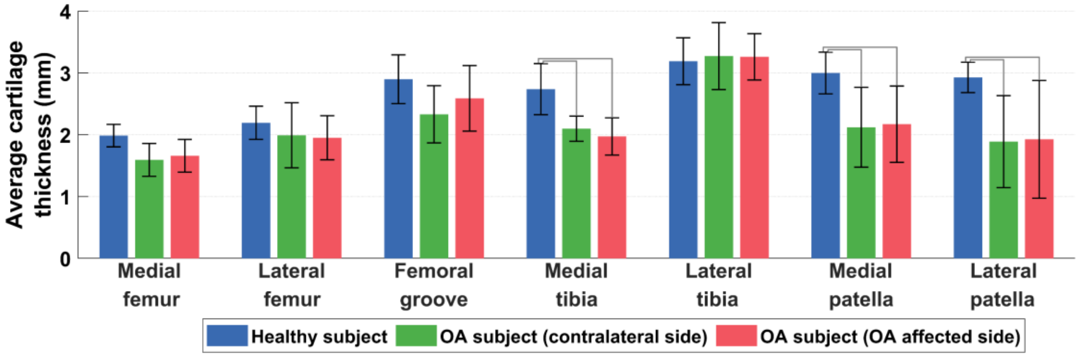


**Fig. S5.** Knee cartilage thickness from the fourth dataset of the study, using proton density (COR-IW-TSE approximate) sequence. Gray lines indicate statistically different results ($p<0.05$, Kruskal-Wallis analysis of variance).

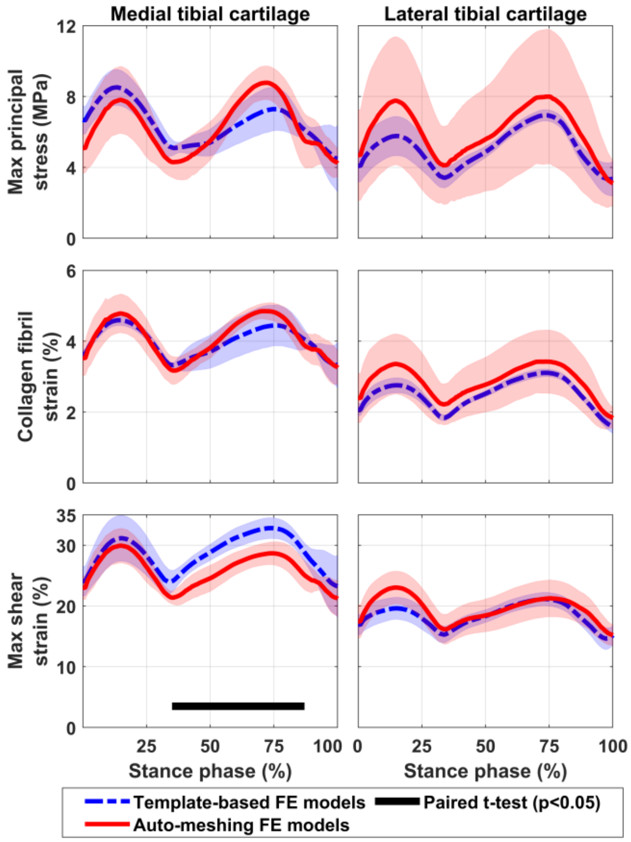


**Fig. S6.** Tissue mechanics within the medial and lateral tibial cartilage estimated by template-based (in blue) and auto-meshing (in red) finite element (FE) models of the study, using proton density (COR-IW-TSE approximate) images of the fourth dataset of the study. The black line shows time points at which results from the two FE modeling methods were statistically different.

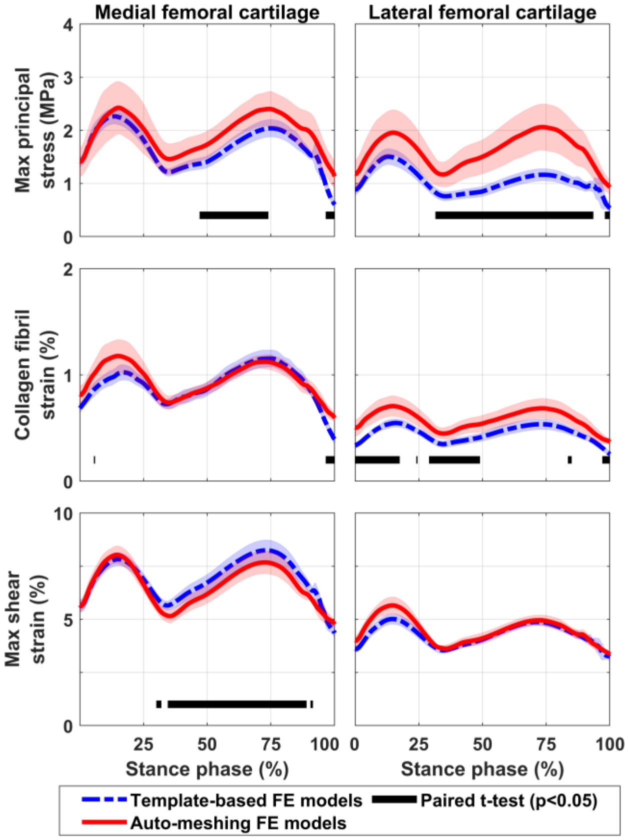


**Fig. S7.** Tissue mechanics within the medial and lateral femoral cartilage estimated by template-based (in blue) and auto-meshing (in red) finite element (FE) models of the study, using proton density (COR-IW-TSE approximate) images of the fourth dataset of the study. The black line shows time points at which results from the two FE modeling methods were statistically different.

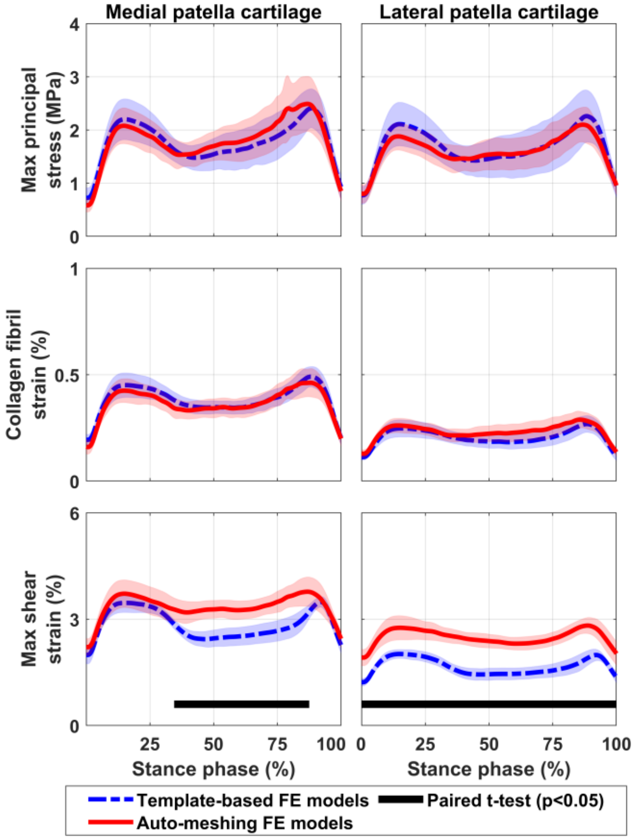


**Fig. S8.** Tissue mechanics within the medial and lateral patellar cartilage estimated by template-based (in blue) and auto-meshing (in red) finite element (FE) models of the study, using proton density (COR-IW-TSE approximate) images of the fourth dataset of the study. The black line shows time points at which results from the two FE modeling methods were statistically different.

[5] R. L. Lenhart, J. Kaiser, C. R. Smith, and D. G. Thelen, "Prediction and validation of load-dependent

behavior of the tibiofemoral and patellofemoral joints during movement," *Ann. Biomed. Eng.*, vol. 43, no. 11, pp. 2675–2685, Nov. 2015, doi: 10.1007/s10439-015-1326-3.

[6] A. Esrafilian *et al.*, "12 Degrees of Freedom Muscle Force Driven Fibril-Reinforced Poroviscoelastic Finite Element Model of the Knee Joint," *IEEE Trans. Neural Syst. Rehabil. Eng.*, vol. 29, pp. 123–133, 2021, doi: 10.1109/TNSRE.2020.3037411.

[7] A. Esrafilian *et al.*, "An EMG-assisted muscle-force driven finite element analysis pipeline to investigate joint- and tissue-level mechanical responses in functional activities: towards a rapid assessment toolbox," *IEEE Trans. Biomed. Eng.*, vol. 69, no. 9, pp. 2860–2871, Sep. 2022, doi: 10.1109/TBME.2022.3156018.

[8] W. Schroeder, K. Martin, and B. Lorensen, *The Visualization Toolkit*, 4th ed. Kitware, 2006.

[19] B. M. Leo, M. A. Turner, and D. R. Diduch, "Split-line pattern and histologic analysis of a human

osteochondral plug graft," *Arthrosc. J. Arthrosc. Relat. Surg.*, vol. 20, pp. 39–45, Jul. 2004, doi: 10.1016/J.ARTHRO.2004.04.043.

10.1177/1947603510368689.
